# GOLDEN TANGERINE TOMATO ROOTS SHOW INCREASED ACCUMULATION OF ACYCLIC CAROTENOIDS, LESS ABSCISIC ACID, DROUGHT SENSITIVITY, AND IMPAIRED ENDOMYCORRHIZAL COLONIZATION

**DOI:** 10.1101/2021.12.07.471680

**Authors:** Jwalit J Nayak, Sidra Anwar, Priti Krishna, Zhong-Hua Chen, Jonathan M. Plett, Eloise Foo, Christopher I. Cazzonelli

**Author notes:** **Corresponding Author:** Christopher I Cazzonelli. The author responsible for the distribution of materials integral to the findings presented in this article is: Christopher Cazzonelli. Equal authorship (Jwalit J Nayak, Sidra Anwar).

## Abstract

Heirloom golden tomato fruit varieties are highly nutritious as they accumulate tetra-*cis*-lycopene, which has a higher bioavailability and recognised health benefits in treating anti-inflammatory diseases compared to all-*trans*-lycopene isomers found in red tomatoes. We investigated if photoisomerization of tetra-*cis*-lycopene occurs in roots of the golden *tangerine* Micro-Tom variety (*tang*^*mic*^), and how this affects root to shoot biomass, mycorrhizal colonization, abscisic acid accumulation, and responses to drought. *tang*^*mic*^ plants grown in soil under glasshouse conditions displayed a reduction in height, number of flowers, fruit yield, and root length compared to wild type (WT). Soil inoculation with *Rhizophagus irregularis* revealed fewer arbuscules and other fungal structures in the endodermal cells of roots in *tang*^*mic*^ relative to WT. The roots of *tang*^*mic*^ hyperaccumulated acyclic *cis*-carotenes, while only trace levels of xanthophylls and abscisic acid were detected. In response to a water deficit, leaves from the *tang*^*mic*^ plants displayed a rapid decline in maximum quantum yield of photosystem II compared to WT, indicating a defective root to shoot signalling response to drought. The lack of xanthophylls biosynthesis in *tang*^*mic*^ roots reduced abscisic acid levels, thereby likely impairing endomycorrhiza colonisation and drought-induced root to shoot signalling.

**Research Highlights:**

1. Photoisomerization of prolycopene to lycopene is limited in root plastids.
2. Roots of *tangerine* reveal an important tissue sink to store micronutrients such as prolycopene.
3. Roots of *tangerine* lack ABA and show impaired mycorrhizal colonization.
4. The *tangerine* plant is drought sensitive and has a smaller biomass as well as reduced yield.

## INTRODUCTION

Consumption of carotenoid-rich fruits and vegetables has been associated with a decreased risk of some cancers, cardiovascular diseases, macular degeneration, and cataract formation (Olson, 1999). Lycopene provides tomatoes with a reddish coloured pigmentation, and is primarily consumed through eating raw tomatoes and/or tomato-based products (Canene-Adams *et al*., 2005; Clinton, 1998). The thermodynamically stable all-*trans* configuration of lycopene constitutes 95% of lycopene forms found in red tomatoes (Porrini *et al*., 1998). In human blood plasma and tissues, 50 % of total lycopene constitutes *cis*-isomers, with all-*trans*-, 5-*cis*-, 9-*cis*-, 13-*cis*-, and 15-*cis*-lycopene being common types (Clinton *et al*., 1996). The heirloom *tangerine* (*tang*^*mic*^) tomato fruit varieties accumulate tetra-*cis*-lycopene (pro-lycopene), along with phytoene, phytofluene, ζ-carotene, and neurosporene relative to other tomato varieties, giving its fruit a characteristic orange to golden colour (Tanambell *et al*., 2020). Consumption of *tangerine* tomato fruits leads to an increase in the absorption of the *cis* isoforms of lycopene and higher bioavailability, offering better potential for treating or preventing cancer and other anti-inflammatory diseases (Tanambell *et al*., 2019; Unlu *et al*., 2007).

Carotenoid biosynthesis in plants involves two distinct parts; firstly, a linear pathway that evolved to produce several acyclic carotenes (e.g. phytoene, phytofluene, di-*cis*-ζ-carotene, tri-*cis*-ζ-carotene, neurosporene and tetra-*cis*-lycopene, all-*trans* lycopene) that are barely detectable in most tissues (Alagoz *et al*., 2018), and a second pathway that diverges into epsilon-beta branches to generate the more abundant cyclic carotenoids (e.g. lutein, β-carotene, violaxanthin and neoxanthin) (Baranski and Cazzonelli, 2016). Genetic perturbations that impair the function of the CAROTENOID ISOMERASE (CRTISO) rate-limit the isomerization of tetra-*cis*-lycopene (prolycopene) into all-*trans* lycopene, leading to the accumulation of acyclic *cis*-carotenes in the absence of light (Alagoz *et al*., 2018). Light can mediate the *cis* to *trans* switch in carotenoid configuration of carbon bonds in the presence of a photosensitiser, allowing photoisomerization of prolycopene to all-*trans*-lycopene to proceed in photosynthetic leaf tissues (Alagoz *et al*., 2018; Anwar *et al*., 2021; Isaacson *et al*., 2002; Vijayalakshmi *et al*., 2015). In young emerging leaves from Arabidopsis *crtiso* mutant plants growing under shorter photoperiod, a *cis*-carotene derived apocarotenoid signal (ACS) impairs chloroplast biogenesis (Cazzonelli *et al*., 2020) and was attributed to leaf virescent phenotypes (yellowing of leaves) reported in maize, melon, rice, and tomato mutants that lack CRTISO activity (Alagoz *et al*., 2020; Chen *et al*., 2010; Galpaz *et al*., 2013; Isaacson *et al*., 2002; Li *et al*., 2007; Park *et al*., 2002). In contrast, chromoplasts in fruit tissues from the MicroTom *tangerine* mutant variety (*tang*^*mic*^) which lack CRTISO activity, cannot photoisomerize prolycopene to all-*trans*-lycopene despite sufficient light penetration (Isaacson *et al*., 2002). What remains unknown is whether or not leucoplasts from root tissues can photoisomerize prolycopene.

Enzymatic and non-enzymatic cleavage of carotenoids result in apocarotenoid signalling metabolites that function to control plastid biogenesis and gene expression, mediate plant development, and modulate root-mycorrhizal interactions (Cazzonelli *et al*., 2020; Escobar-Tovar *et al*., 2021; Fiorilli *et al*., 2019; Moreno *et al*., 2021; Walter *et al*., 2010). Carotenoids (e.g. β-carotene, zeaxanthin, violaxanthin and neoxanthin) provide precursors for the biosynthesis of two key phytohormones, abscisic acid (ABA) and strigolactone (SL), that mediate stress responses and control shoot bud outgrowth, respectively (Al-Babili and Bouwmeester, 2015; Finkelstein, 2013; Jia *et al*., 2018). Mutations that impair acyclic *cis*-carotene biosynthesis (e.g. prevent/reduce xanthophyll biosynthesis) have been shown to increase shoot branching in Arabidopsis presumably by altering SL biosynthesis (Cazzonelli *et al*., 2009), as well as produce ABA-deficient phenotypes associated with precocious seed germination (viviparous), reduced seed dormancy, and compromised stress responses in Arabidopsis, maize and rice (Fang *et al*., 2008; McCarty, 1995). Physiological responses of tomatoes to drought stress reveal that biomass production is tightly linked to stomatal closure and increased ABA biosynthesis (Blum, 2009; Xiong *et al*., 2006). The production of ABA in the roots and its transport to the leaves provides a mechanism for the plant to signal soil water status to foliar tissues in order to retain water content and avoid photooxidative stress to the photosystems (Schachtman and Goodger, 2008; Sharp *et al*., 1994). Changes in photosynthetic parameters during water deficit conditions can be monitored using chlorophyll fluorescence imaging and the maximum efficiency of photosystem II (F_v_/F_m_) as a measure of survival (Woo *et al*., 2008). The impact that the loss of CRTISO activity in *tang*^*mic*^ might have on root morphology, ABA accumulation and drought responses remains unknown and yet it could impact upon crop physiology to adversely affect yield.

ABA, SL, zeaxanthin-derived zaxinones, blumenols, and mycorradicin have been shown to play important roles in mediating mycorrhizal symbioses (Fiorilli *et al*., 2019; Hill *et al*., 2018; Hill *et al*., 2021; Walter *et al*., 2010). ABA has a positive and direct role in root colonisation by arbuscular mycorrhizal fungi (AMF) (Charpentier *et al*., 2014). For example, the ABA-deficient tomato mutant, *sitiens*, showed a reduction in AMF colonisation, arbuscular formation, and impaired functionality due to increased ethylene levels (Herrera□Medina *et al*., 2007; Martín□Rodríguez *et al*., 2011). Exogenous application of ABA at low concentrations in *Medicago trunculata* promoted AMF fungal colonisation (Charpentier *et al*., 2014). SL can also enhance AMF colonisation by promoting spore germination and hyphal branching of the AMF (López-Ráez *et al*., 2017). AMF help to facilitate soil mineralisation, nutrient uptake (e.g. phosphorus) and protect their host in natural and cultivated soils by improving tolerance to biotic and abiotic stresses such as drought (Mitra *et al*., 2021; Rouphael *et al*., 2015). It remains unclear if roots of *tang*^*mic*^ mutants can accumulate xanthophylls and if there is any impairment in AMF colonisation.

We hypothesised that the roots from *tang*^*mic*^ plants growing in soil within a glasshouse environment would accumulate acyclic *cis-*carotenes due to impaired photoisomerization, leading to a reduction in xanthophylls, ABA levels, and early sensitivity to drought. We investigated the symbiotic relationship between soil-grown roots of *tang*^*mic*^ plants and the AMF species *Rhizophagus irregularis*. We quantified crop growth and yield, to decipher which horticultural cropping traits (e.g. drought tolerance) might be impacted within the golden *tang*^*mic*^ variety. This study helps to facilitate knowledge regarding the protected cropping of golden tomato fruit varieties that offer superior nutritional and health benefits through the enriched accumulation of prolycopene.

## MATERIALS AND METHODS

### Experimental materials and growth conditions

The tomato seeds from each germplasm were initially grown in 20 cell plastic trays (Garden City Plastics, Dandenong South, VIC, Australia) for 14 days and then transplanted to 200mm pots carrying approximately 4.5 litres of soil, with two plants in each pot. The transplanted seedlings were grown in a blend of Tomato and Vegetable growing mix and Osmocote slow-release fertiliser (Garden City Plastics) 16-3.5-9.1 (N-P-K) (30 g/5L) under a glasshouse facility in Richmond, NSW 2753. The entire experiment was conducted twice at an interval of 8 weeks between August and November. The glasshouse temperature during daytime was set to 23°C ± 2°C and 17°C ± 2°C for nights. Plants were watered at regular intervals of three times per week. Phenotypic observations such as fresh weights (FW), dry weights (DW), plant height, primary root length, number of branches, fruits and flowers were carried out 8-weeks after germination.

### Media Preparation

Seeds of WT and *tang*^*mic*^ were sterilised using chlorine gas in a sealed container for 3 hours, followed by washing seeds once with 70% ethanol and three times with sterilised water. Seeds were plated on Murashige and Skoog (MS) media (Caisson, MSP01) supplemented with Gamborg’s half-strength vitamin solutions (0.5 mL/L) (Sigma Aldrich), 0.5% phytagel (Sigma-Aldrich) (5 g/L), and 1 % sucrose in 120 ×120 × 17 mm square Petri dishes. Seeds were grown vertically under a 16-hour photoperiod at 22°C, 60% relative humidity and light intensity of 120 μmol m^-2^ s^-1^. Tomato phenotypes were analysed on day 14 using ImageJ software.

### Pigment analysis

Five individual plants from WT and *tang*^*mic*^ were selected under the glasshouse facility to study the carotenoid accumulation in leaves and roots. Two to three mature green leaves were cut from the same branch with a scissor and snap-frozen after determining the fresh weight (∼100 mg). Roots were cut into two halves, weighed and frozen in liquid nitrogen separately. One part was used for pigment extraction (∼ 150 mg), while the other half was used for phytohormone quantification. Frozen leaf tissues were homogenised to a fine powder using TissueLyser® (QIAGEN) while root tissues, on the other hand, were homogenised using mortar and pestle. Carotenoids were extracted as previously described (Alagoz *et al*., 2020) using a reverse-phase high-performance liquid chromatography (HPLC) (Agilent 1260 Infinity) and a YMC-C30 (250 × 4.6 mm, S-5μm) column. Carotenoids and chlorophylls were identified based upon their retention time relative to known standards and their light emission absorbance spectra at 440 nm (chlorophyll, β-carotene, xanthophylls, pro-neurosporene, tetra-*cis*-lycopene, other neurosporene and lycopene isomers), 400 nm (ζ-carotenes), 340 nm (phytofluene) and 286 nm (phytoene). Absolute quantification of xanthophyll pigments was performed as described (Anwar *et al*., 2022).

### Quantification of ABA in roots

For quantifying endogenous hormones, five replicates of 8-weeks old root tissues from each genotype were harvested (∼200 mg), snap-frozen and stored at -80°C. Hormone extraction was performed previously described (Brenya *et al*., 2020; Hill *et al*., 2021) and analysed using the UPLC/ESI-MS/MS (Water, Milford, USA). The external standard of ABA was obtained from Sigma Aldrich. The internal standard, a deuterated compound of [^2^H_6_](+)-*cis,trans*-ABA, was obtained from Olchemim Ltd, Olomouc, Czech Republic. For the analysis of the extracts, a HALO™ C18 (Advanced Materials Technology, Inc., Wilmington, USA) column (2.1 × 75 mm, 2.7 μm) was used. Calibration curves for each analyte were generated using MassLynx 4.1™ software (Waters, USA).

### Drought stress and relative water content

MicroTom WT and *tang*^*mic*^ plants were grown in seed and cutting soil mixture (DEBCO Pty, Australia) with Osmocote fertiliser in a growth chamber at 22°C and illuminated by approximately 190 μmol m^-2^ s^-1^ under 16h photoperiod. All plants were watered for the first three weeks, and each germplasm was divided into two groups, i.e., well-watered (WW), acting as controls (n=2) and drought-stressed (DS) (n=8) for each germplasm. For analysis, the plants were dark-adapted for 20min in a dark room. The F_v_/F_m_ was measured with a PAM2500 (Walz, Effeltrich, Germany) operated with the PamWin_3 software. The measurements were taken starting from Day 0, followed by every consecutive day until the plants dried out.

For relative water content, measurements were taken every five days. Under WW and DS conditions, two-three fully expanded leaves from each germplasm were selected. Weights were taken as per (Zhou *et al*., 2017) with little modification. After cutting, the FW was immediately measured and then the leaves were immersed in ddH_2_O in a petri dish and incubated at room temperature. After four hours, the leaves were taken out, wiped to remove surface water, and weighed to obtain turgid weights (TW). Following that, the leaves were dried in an oven at 80°C for 48 hours and weighed to obtain DW. The formula used for estimating the percentage of relative water content (RWC; defined as [(FW-DW)/(TW-DW)] × 100.)

### Mycorrhizal colonisation

Inoculum for mycorrhizal experiments was live corn pot culture originally inoculated with spores of *Rhizophagus irregularis* (INOQ Advantage, INOQ GMBH, Germany), grown under glasshouse conditions that received modified Long Ashton nutrient solution containing 3.7mM KNO_3_ and 0.05mM NaH_2_PO_4_ once a week. The inoculum contained colonised root segments, external hyphae and spores. Tomato seeds were germinated in potting mix and transplanted two weeks after sowing into 2L pots containing a 2 vermiculite : 2 gravel : 1 inoculum topped with 2cm of vermiculite. Plants were grown under short-day photoperiod (natural daylight for 8 hours and then transferred to dark chamber). Plants were supplied with 75ml/pot modified Long Ashton nutrient solutions (Hewitt, 1966) containing 2.5 mM KNO_3_ and mM NaH_2_PO_4_ twice a week. Plants were harvested six weeks after transplanting. Fresh shoot and root weights was recorded, and then root segments of approx. 1.5 cm in length were taken and stored in 50% ethanol until staining. Root segments were stained for mycorrhizal structures using the ink and vinegar method (Vierheilig *et al*., 1998). Mycorrhizal colonisation of roots was scored as described previously (McGonigle *et al*., 1990), where 150 intersects were observed from 25 root segments per plant. The presence of arbuscules, vesicles and intraradical hyphae at each intersects were scored separately. The total colonisation of mycorrhizae was calculated as the percentage of intersects that have the presence of any fungal structures, and arbuscular frequency was calculated from the percentage of intersects that contained arbuscules.

### Statistical analysis

Analysis of Variance (ANOVA) was used to test for significant differences between the treatments using SigmaPlot 14.0 software.

## RESULTS

### *tang*^*mic*^ shows reduced growth, shoot biomass, fruit yield and altered root architecture

The plant height, number of shoot branches, flowers and fruits were assessed in WT and *tang*^*mic*^ plants. The newly emerged leaves of *tang*^*mic*^ displayed a virescent yellow leaf phenotype, consistent with previous reports (Fig. 1A) (Cazzonelli *et al*., 2020; Isaacson *et al*., 2002). The *tang*^*mic*^ plants showed a significant reduction in plant height (1.3-fold), the number of flowers (3.6-fold) and fruits (1.9-fold), and a significant increase in shoot branching (1.3-fold) compared to WT (Fig. 1B-E). Consistent with previous observation (Isaacson *et al*., 2002), the fruits of *tang*^*mic*^ appeared golden orange in colour compared to the red-pigmented colour of WT (Fig. 1F). The shoot biomass of *tang*^*mic*^ was 2-fold lower in terms of both dry and fresh weight than WT (Fig. 1G). The fresh weight of *tang*^*mic*^ roots was slightly higher, while the dry weight was not significantly different from WT. The dry root to shoot ratio of biomass was similar between WT and *tang*^*mic*^ (Fig. 1H). The *tang*^*mic*^ plants displayed an overall reduction in size compared to WT (Fig. 1I). The root phenotypes (primary root length, number of lateral roots, and root thickness) of *tang*^*mic*^ were assessed by growing plants on artificial media (MS) under 16 h of fluorescent illumination for 14 days (Fig. 2). *tang*^*mic*^ plants grown on media showed a significant decrease (2-fold) in primary root length compared to WT (Fig. 2B). A similar reduction in primary root length was observed in seedlings grown in soil for 8-weeks within a glasshouse (Fig. 2A-C and Fig. 1I). The number of lateral roots was severely reduced in *tang*^*mic*^ (4-fold), yet there was a slight increase (1.3-fold) in root thickness as compared to WT (Fig. 2D-E).

**Figure 1:**
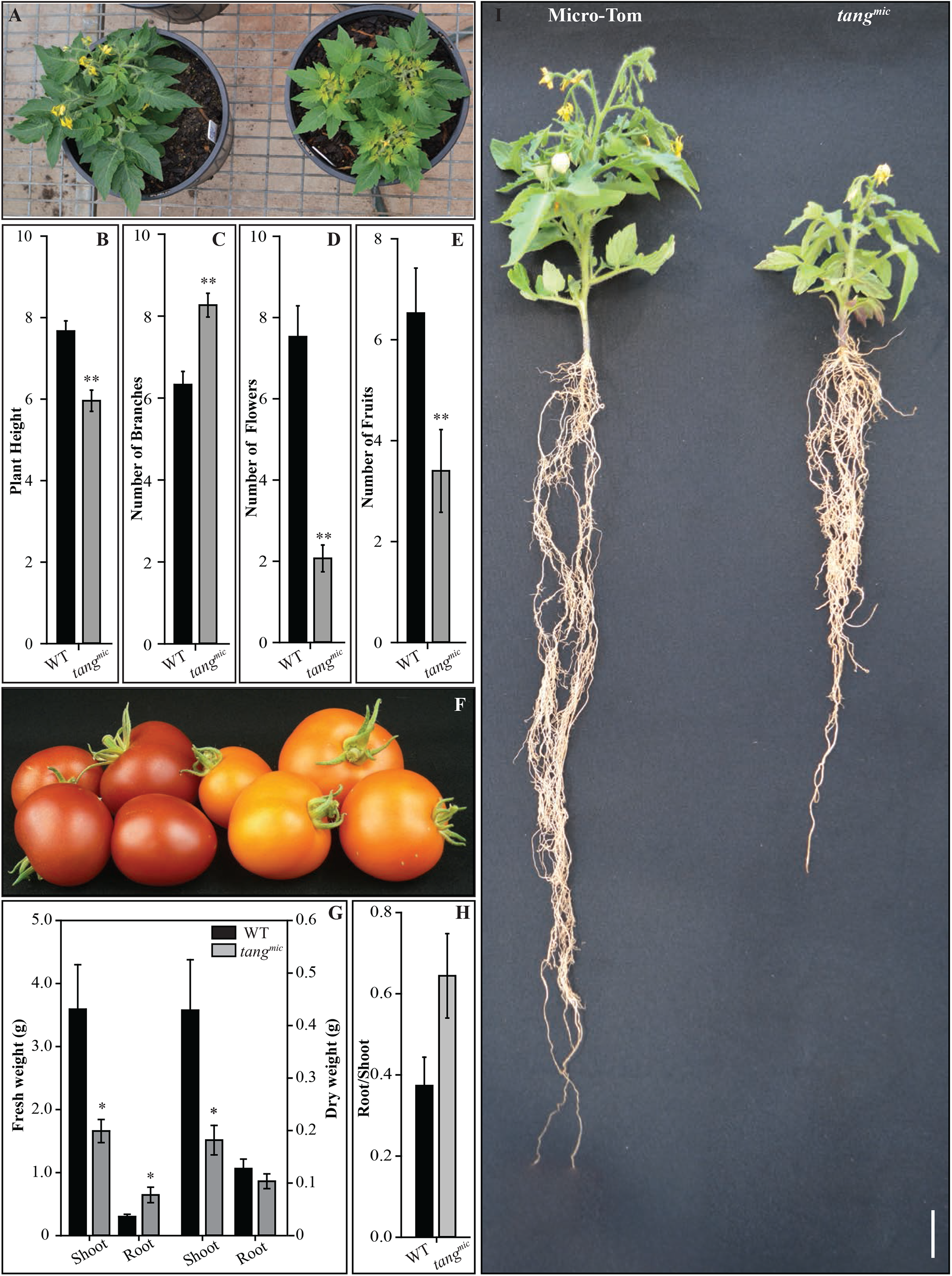
Phenotypic comparison of Micro-Tom (WT) and *tang*^*mic*^ (tangerine) plants grown in soil within a glasshouse. Plants n in soil for 8 weeks in the glasshouse under the Australian spring/summer photoperiod and natural solar **(A)** Representative phenotypes of *tang*^*mic*^ showing a yellow virescence in newly emerged leaves. Fifteen plants were selected for each germplasm and **(B)** plant height (cm), **(C)** number of branches (cm), **(D)**, number of fruits (cm) **(E)**, and number of flowers (cm) were quantified. **(F)** Red (WT) and orange coloured (*tang*^*mic*^) fruits, **(G)** fresh and dry weights and root biomass and **(H)** ratio of roots to shoots are shown. **(I)** The significant difference was determined by one-way ANOVA. The p-values (*0.05; **0.01) and standard error (n=15) bars are displayed. The scale bar represents 1cm.

**Figure 2:**
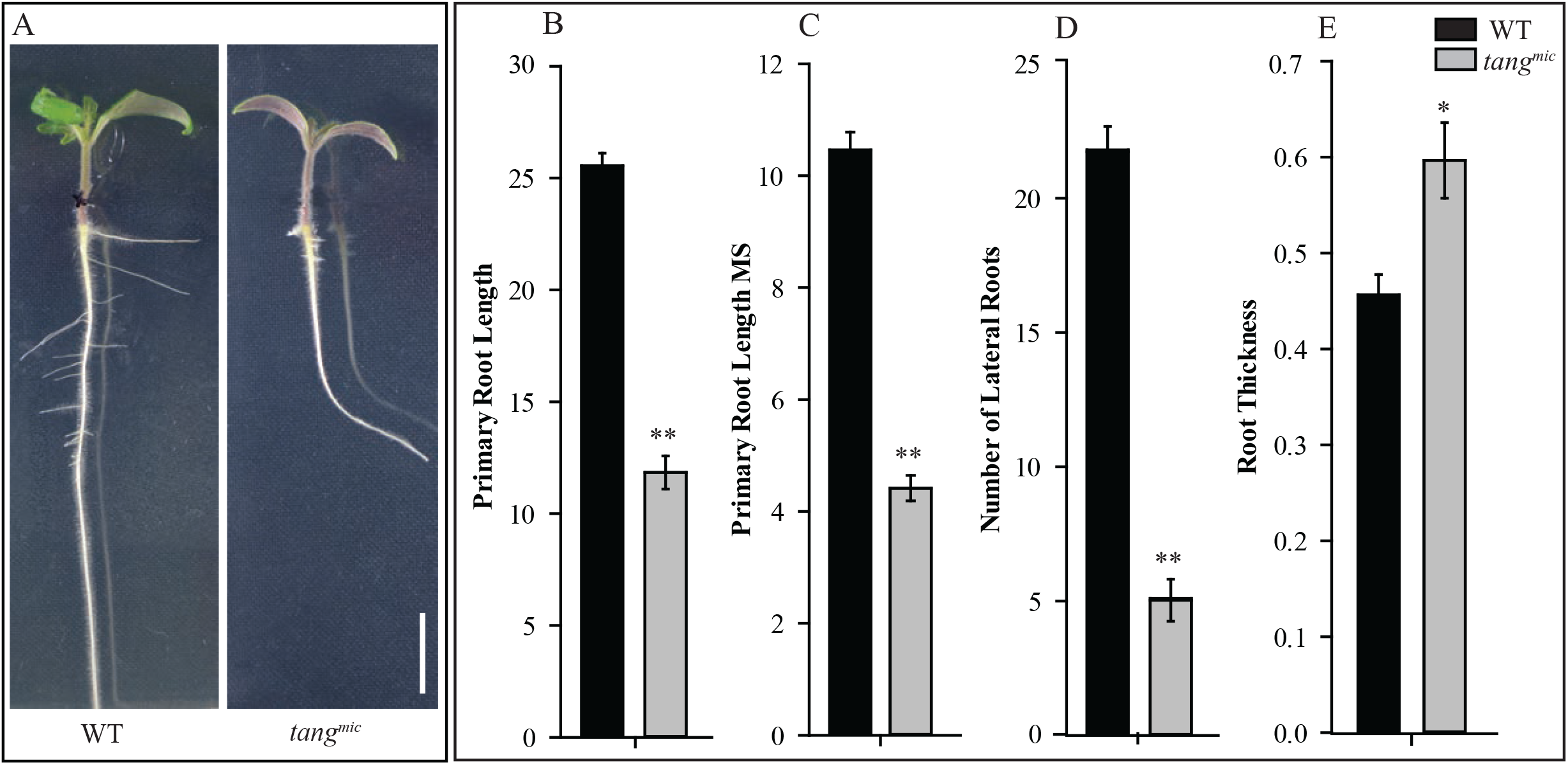
Comparison of root phenotypes displayed by WT and *tang*^*mic*^ grown in soil and artificial media. **(A)** Representative picture of WT and *tang*^*mic*^ plants grown on artificial (MS) media in an environmentally controlled growth chamber (120 μmol m^-2^ s^-1^ of light and 16 hr photoperiod). **(B)** Average primary root length (cm) of 8-week-old *tang*^*mic*^ plants grown in soil under glasshouse conditions (n=15). **(C)** Average length (cm) of the primary root nd *tang*^*mic*^ seedlings grown on MS media. **(D)** Average number of lateral roots displayed by WT and dlings on MS media. **(E)** Average root thickness (cm) showed by WT and *tang*^*mic*^ plants on MS media (Zoom %= 600). Bars represent the standard error of the mean (n=10). Asterisk represents significant differences between and *tang*^*mic*^ roots (one-way ANOVA; p-values *0.05; **0.01). The scale bar represents 1cm.

### *tang*^*mic*^ roots hyperaccumulate acyclic cis-carotenes and lack xanthophylls

Absolute levels of carotenoids and their composition were quantified in mature leaves and roots from *tang*^*mic*^ and WT (Fig. 3; Table 1). An overlay in HPLC chromatograms from WT and *tang*^*mic*^ revealed a similar carotenoid profile in mature leaves, except for one β-carotene isomer being more abundant in *tang*^*mic*^ (Fig. 3B). Mature leaves from *tang*^*mic*^ showed a significant reduction in neoxanthin (3-fold) and lutein (4-fold), while violaxanthin and β- carotene remained similar, leading to an overall 1.5-fold lower total carotenoid content compared to WT (Fig. 3A). Accordingly, the percent carotenoid composition relative to the total pool for neoxanthin and lutein was lower in *tang*^*mic*^ leaves, while those of violaxanthin and β-carotene were significantly higher (Table 1). The total absolute chlorophyll levels remained unchanged in both genotypes, even though there was a small significant reduction in Chl b in *tang*^*mic*^, which significantly enhanced the chlorophyll a/b ratio relative to WT (Fig. 3A; Table 1).

**Table 1:**
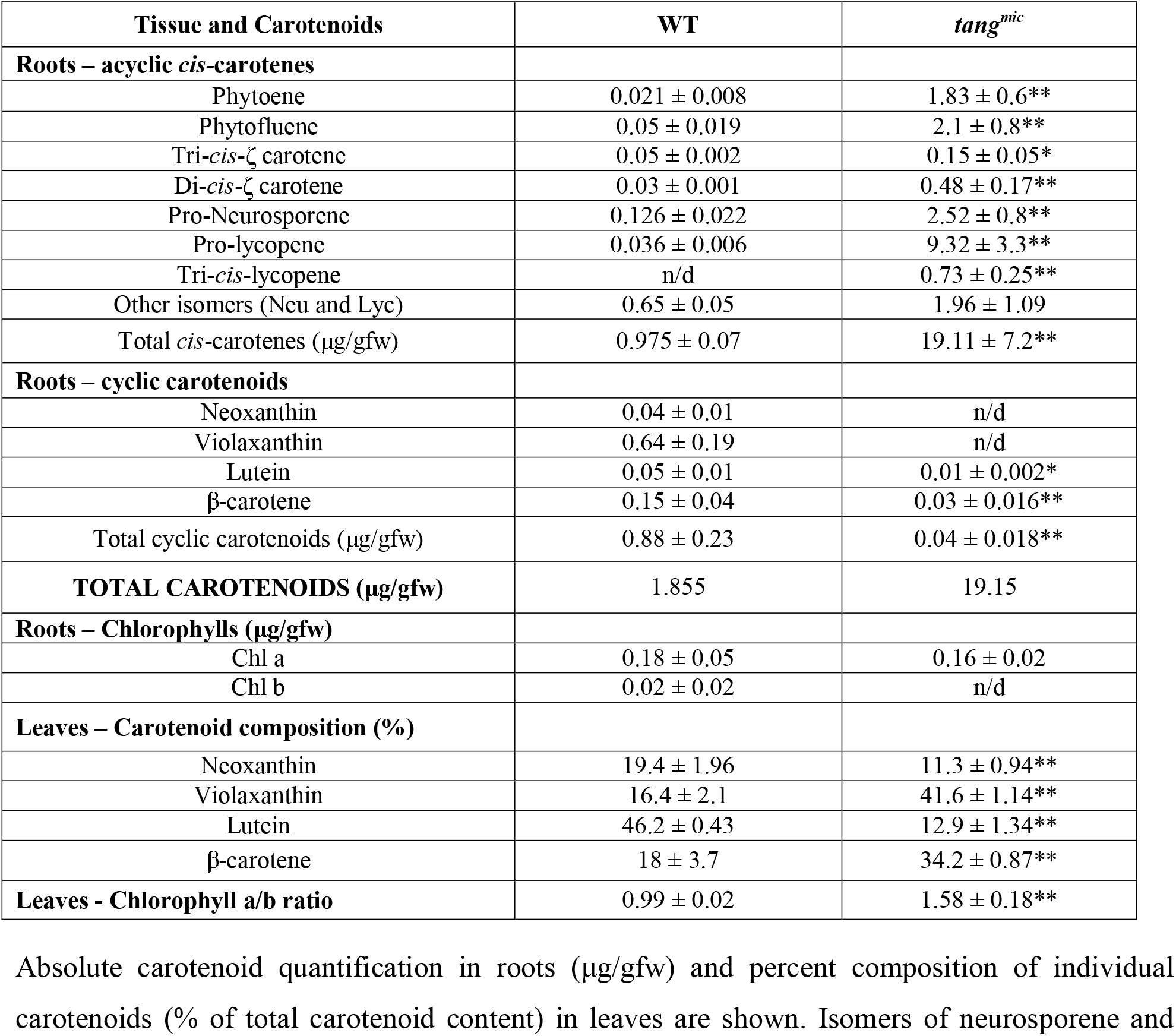
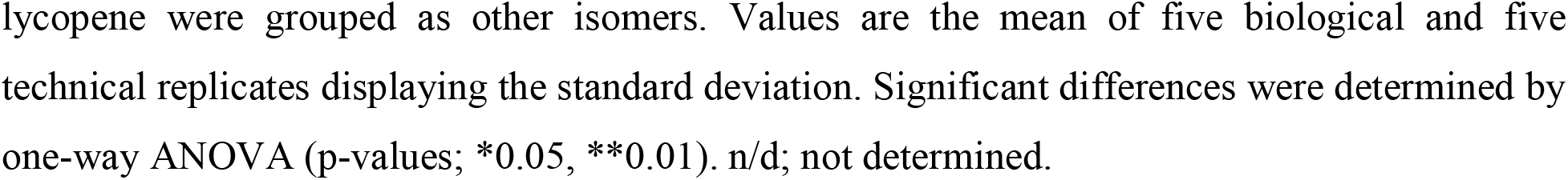
Carotenoid levels and composition in roots and leaves of WT and *tang*^*mic*^ plants grown within a glasshouse.

| Tissue and Carotenoids | WT | *tang<sup>mic</sup>* |
| --- | --- | --- |
| **Roots – acyclic *cis*-carotenes** |  |  |
| Phytoene | 0.021 ± 0.008 | 1.83 ± 0.6** |
| Phytofluene | 0.05 ± 0.019 | 2.1 ± 0.8** |
| Tri-*cis*-ζ carotene | 0.05 ± 0.002 | 0.15 ± 0.05* |
| Di-*cis*-ζ carotene | 0.03 ± 0.001 | 0.48 ± 0.17** |
| Pro-Neurosporene | 0.126 ± 0.022 | 2.52 ± 0.8** |
| Pro-lycopene | 0.036 ± 0.006 | 9.32 ± 3.3** |
| Tri-*cis*-lycopene | n/d | 0.73 ± 0.25** |
| Other isomers (Neu and Lyc) | 0.65 ± 0.05 | 1.96 ± 1.09 |
| Total *cis*-carotenes (μg/gfw) | 0.975 ± 0.07 | 19.11 ± 7.2** |
| **Roots – cyclic carotenoids** |  |  |
| Neoxanthin | 0.04 ± 0.01 | n/d |
| Violaxanthin | 0.64 ± 0.19 | n/d |
| Lutein | 0.05 ± 0.01 | 0.01 ± 0.002* |
| β-carotene | 0.15 ± 0.04 | 0.03 ± 0.016** |
| Total cyclic carotenoids (μg/gfw) | 0.88 ± 0.23 | 0.04 ± 0.018** |
| **TOTAL CAROTENOIDS (μg/gfw)** | 1.855 | 19.15 |
| **Roots – Chlorophylls (μg/gfw)** |  |  |
| Chl a | 0.18 ± 0.05 | 0.16 ± 0.02 |
| Chl b | 0.02 ± 0.02 | n/d |
| **Leaves – Carotenoid composition (%)** |  |  |
| Neoxanthin | 19.4 ± 1.96 | 11.3 ± 0.94** |
| Violaxanthin | 16.4 ± 2.1 | 41.6 ± 1.14** |
| Lutein | 46.2 ± 0.43 | 12.9 ± 1.34** |
| β-carotene | 18 ± 3.7 | 34.2 ± 0.87** |
| **Leaves - Chlorophyll a/b ratio** | 0.99 ± 0.02 | 1.58 ± 0.18** |

Absolute carotenoid quantification in roots (μg/gfw) and percent composition of individual carotenoids (% of total carotenoid content) in leaves are shown. Isomers of neurosporene and
lycopene were grouped as other isomers. Values are the mean of five biological and five technical replicates displaying the standard deviation. Significant differences were determined by one-way ANOVA (p-values; \*0.05, \*\*0.01). n/d; not determined.

**Figure 3:**
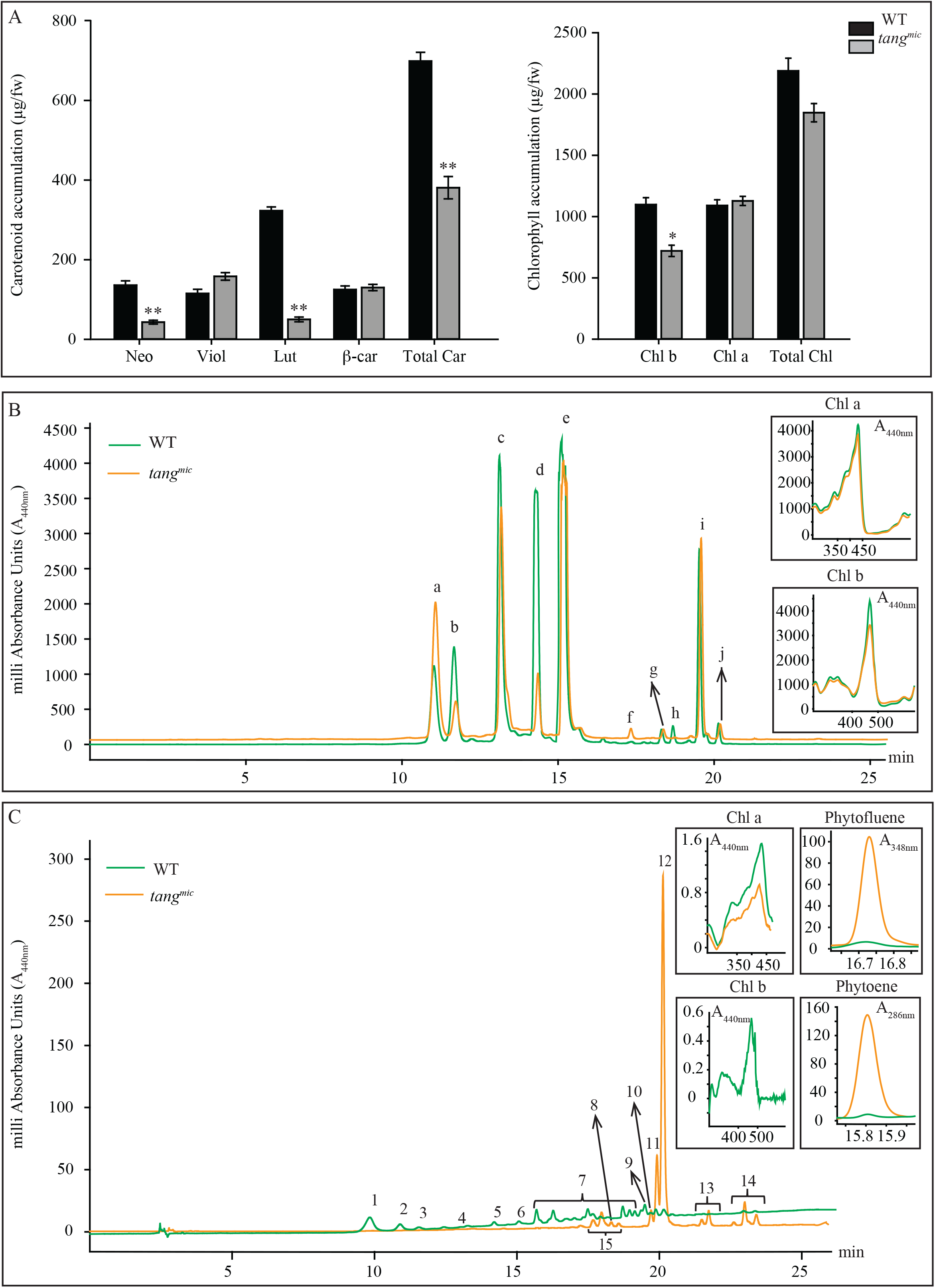
Carotenoid accumulation in mature leaves and roots of WT and *tang*^*mic*^ plants grown in soil within a glasshouse. **(A)** Carotenoid and chlorophyll levels (μg/gfw) in mature leaf tissues (n=5). Carotenoid profiles of WT and *tang*^*mic*^, leaf (**B**)and root (**C**) tissues depicted by a representative HPLC chromatogram traces. The y-axis value represents the milli absorbance units (mAU) and x-axis is the retention time of carotenoid detection. Carotenoids identified for leaves are labelled as: violaxanthin (a), neoxanthin (b), Chl *b* (c) lutein (d), Chl *a* (e), β-carotene isomer-1 (f), e cis-isomer 1 (g); β-carotene isomer-1 (h) β-carotene (i); β-carotene isomer-2 (j). Carotenoids identified for labelled as: 9-cis-violaxanthin (1), violaxanthin (2), neoxanthin (3), Chlorophyll-b (4), lutein (5), Chlorophyll-a (6), neurosporene isomers (7), tri-cis-ζcarotene (8), β-carotene (9), di-cis-ζcarotene (10), pro-neurosporene (11), pro-lycopene (12), neurosporene isomers (13), lycopene isomers (14), and neurosporene 15). Inserts show absorbance spectra for phytoene (A_286nm_), phytofluene (A_348nm_), Chl a(A_440nm_) and Chl b (A_440nm_). Data are representative of multiple samples from at least two experimental repetitions. Statistical analysis performed using a t-test (p-value *0.05; ** 0.01).

The absolute carotenoid and chlorophyll levels in roots of soil-grown plants were strikingly different between the genotypes (Fig. 3C; Table 1). WT roots accumulated β-carotene, neoxanthin, violaxanthin, and lutein (0.88 μg/gfw), which were either trace (β-carotene, lutein) or undetected (neoxanthin, violaxanthin) in *tang*^*mic*^ roots (0.04 μg/gfw). This indicates a block in xanthophyll biosynthesis (Table 1). WT roots accumulated trace levels of acyclic *cis-*carotenes (0.97 μg/gfw), except for tri-*cis*-lycopene that could not be detected. In contrast, soil-grown *tang*^*mic*^ roots showed a significantly higher level of total *cis*-carotenes (19.11 μg/gfw), that was 10.3-fold higher than all carotenoids quantified in WT roots (Table 1). Prolycopene (9.32 μg/gfw) accounted for almost 50% of the total *cis*-carotene pool in *tang*^*mic*^ roots (Table 1). Chl a (0.16 μg/gfw), but not Chl b, was detected in roots of *tang*^*mic*^. In contrast, 0.18 μg/gfw of Chl a and trace levels of Chl b were detected in WT roots, indicating that light could filter through the soil to promote root chloroplasts (Fig. 3C; Table 1).

### *tang*^*mic*^ roots lack ABA and mycorrhizal structures

ABA was quantified in soil-grown roots of WT and *tang*^*mic*^ using mass-spectrometry. The ABA levels were significantly reduced (23-fold) in *tang*^*mic*^ roots compared to WT (Fig. 4A). The reduction in the ABA levels of *tang*^*mic*^ roots was consistent with the lack of detection of ABA precursors, neoxanthin and violaxanthin (Table 1). The percentage colonisation rate of *R. irregularis* was determined by quantifying mycorrhizal arbuscules and other structures (e.g., mycorrhizal hyphae and vesicles) in *tang*^*mic*^ and WT roots. The percent colonisation rate was 30- fold reduced in arbuscules and other structures in *tang*^*mic*^ compared to WT (Fig. 4B). Therefore, *tang*^*mic*^ roots lack ABA and barely have any visible mycorrhizal arbuscules or other structures.

**Figure 4:**
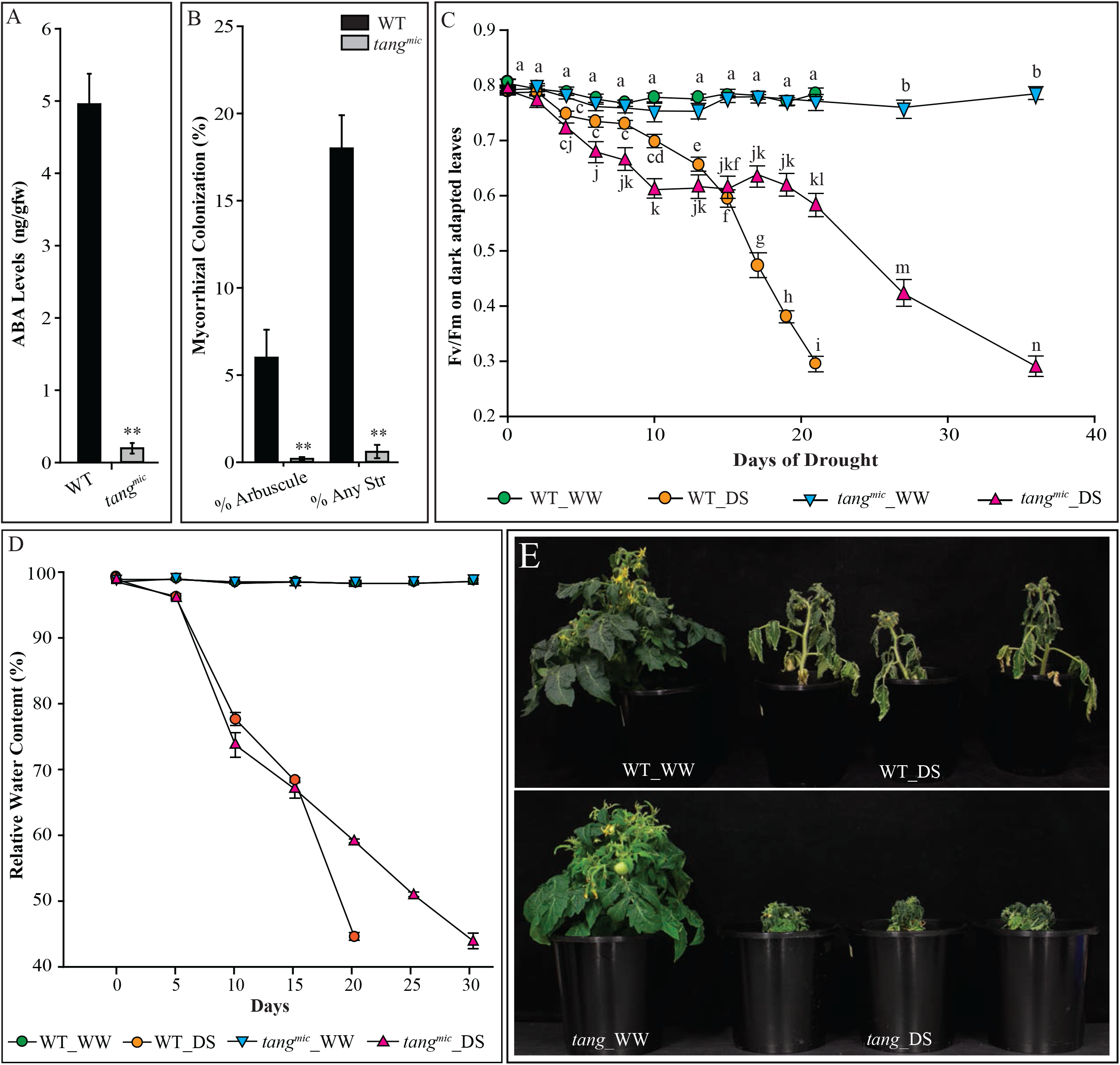
Quantification of endogenous ABA levels from soil-grown WT and *tang*^*mic*^ roots accompanied colonisation, and sensitivity to drought stress. (A) ABA levels were quantified from 8 weeks soil-grown WT and tang-rror bars show the standard error of the mean (n= 5). Asterisk represents significant differences between WT and *tang*^*mic*^, roots using one-way ANOVA (p < 0.01**). The experiment was repeated twice with similar results. (B) Mycorrhizal colonisation in WT and *tang*^*mic*^ plants grown under a short-day photoperiod (8 hours light). The percentage of arbuscules and other structures was determined using binocular microscopy. Error bars show the standard error of the mean (n = 6). Asterisk represents significant differences in mycorrhizal colonisation between WT and *tang*^*mic*^, roots using one-way ANOVA (p < 0.01**). (C) F_v_/F_m_ was measured at a PAR 180-190 μmol m^-2^ s^-1^ at different days on WT and *tang*^*mic*^, dark adapted leaves. Error bars show the standard error of the mean (n = 8 for DS plants and n=2 for WW for each genotype). Alphabetical letters represent significant differences in the F_v_/F_m_ values between WT and *tang*^*mic*^, plants using two-way ANOVA (p-value < 0.05). (D) Relative water content (%) in leaves of well-watered (WW) (Control) and drought stressed and *tang*^*mic*^, plants at respective days after ceasing to water. (E) Representative images of WT and *tang*^*mic*^, plants at Day 20 and 30, respectively showing a single WW and three representative DS plants.

### *tang*^*mic*^ plants show altered responses to drought stress

We next investigated the impact of drought stress on *tang*^*mic*^ and WT plants by quantifying the maximum efficiency of photosystem II (F_v_/F_m_) on dark-adapted leaves and leaf relative water content (RWC) from glasshouse-grown plants subjected to a 30-day period of water deficit. The F_v_/F_m_ values of drought-stressed WT plants were similar to well-watered controls for the first 10 days of water deficit. Thereafter, between days 10 to 20 there was a significant linear decline in F_v_/F_m_ values from 0.7 to 0.3 in drought stressed WT leaves, while the well-watered WT controls consistently maintained higher F_v_/F_m_ values (0.8) (Fig. 4C). After the 20-day period of drought when WT plants F_v_/F_m_ levels reached 0.3, a sign of unlikely recovery (Woo *et al*., 2008), the drought stressed plants appeared severely wilted and the leaf RWC had declined to 44% relative to the well-watered WT controls (Fig. 4D and E). Leaves from the well-watered *tang*^*mic*^ plants showed F_v_/F_m_ values similar to that of WT (0.8) throughout a 30-day growth period (Fig. 4C). Compared to WT, the F_v_/F_m_ values of leaves from drought stressed *tang*^*mic*^ plants showed a significant rapid reduction in the F_v_/F_m_ values within the first 10 days (∼0.8 to 0.6) that plateaued until day 21, and began a subsequent linear decline reaching an F_v_/F_m_ value of 0.3 at day 37 (Fig. 4C). The leaf RWC of drought stressed *tang*^*mic*^ followed a similar trend to that of WT for the first 15 days, declining to 70% relative to the well-watered *tang*^*mic*^ controls (Fig. 4E). However, *tang*^*mic*^ maintained a slightly higher leaf RWC until day 30 where it reached the WT equivalent of 44% leaf RWC when the F_v_/F_m_ values were approximately 0.3 (Fig. 4C and D). Unexpectedly, drought stressed *tang*^*mic*^ plants appeared less wilted at day 37 compared to drought stressed WT plants at day 20 despite having the same leaf RWC (44%) and F_v_/F_m_ values (0.3) (Fig. 4E). The smaller shoot biomass of *tang*^*mic*^ plants (Fig. 1) may have slowed plant water absorption from the rhizosphere and prolonged the plant’s survival during the water deficit.

## DISCUSSION

### Photoisomerization of prolycopene to tetra-cis-lycopene is limited in underground roots

Photoisomerization compensates for the loss-of-function in CRTISO in photosynthetic foliar tissues of tomato, Arabidopsis, rice, and melon. However, this can be less efficient in plants grown under a shorter photoperiod cycle, evident by varying degrees of leaf virescence due to impaired chloroplast biogenesis (Cazzonelli *et al*., 2020; Galpaz *et al*., 2013; Han *et al*., 2012; Isaacson *et al*., 2002). The newly emerged *tang*^*mic*^ leaves appeared virescent despite the plants being grown under a longer photoperiod during spring/summer. The carotenoid composition and chlorophyll levels in WT and *tang*^*mic*^ mature leaf tissues were consistent with a previous report (Isaacson *et al*., 2002). Even though light can penetrate tomato fruit tissues, the chromoplasts lack a chlorophyll-related photosensitiser required to facilitate the *cis/trans*-photoisomerization of lycopene, resulting in a lack of xanthophylls and an accumulation of *cis*-carotenes in *tangerine* mutants (Isaacson *et al*., 2002; Jensen *et al*., 1982; Vijayalakshmi *et al*., 2015). It was unknown if *cis/trans*-photoisomerization can occur in root plastids such as the leucoplast. Light can penetrate underground through the soil where root-localised photoreceptors can detect light and influence underground positive root phototropism growth (Boccalandro *et al*., 2007; Correll *et al*., 2005; Kiss *et al*., 2001; Warnasooriya *et al*., 2011). Stem-piped light can also promote the movement of signalling molecules from shoot to root resulting in the activation of photosynthesis-associated nuclear genes (*PhANGs*) that enable chloroplast development (Lee *et al*., 2016; Oliveria, 1982), indeed we observed small amounts of both chlorophylls in WT roots. Chl a was detected in *tang*^*mic*^ roots that had trace levels of lutein and β-carotene, indicating the presence of a chlorophyll-related photosensitiser that facilitated *cis/trans*-photoisomerization of prolycopene in the absence of CRTISO activity. However, the roots of *tang*^*mic*^ accumulated substantial levels of *cis*-carotene isoforms, particularly prolycopene that represented >40% of the total carotenoid content. This indicated that the photoisomerization was rate-limiting in generating tetra-*cis*-lycopene, and as a result, neoxanthin and violaxanthin substrates required to produce ABA were lacking in *tang*^*mic*^ roots. Thus, CAROTENOID ISOMERASE activity is essential underground to ensure ABA biosynthesis in root plastids.

The *tang*^*mic*^ roots accumulated 14.8-fold higher total carotenoids (acyclic *cis*-carotenes plus xanthophylls) as compared to WT, revealing an important tissue sink to store health-promoting micronutrients such as prolycopene. This was visually evident by a slightly yellow hue in *tang*^*mic*^ roots compared to WT roots that remained colourless. The compartment storage size of plastids affects carotenoid accumulation by providing a greater sink as reported in *hp-3* and *hp-2* tomato mutants (Cazzonelli *et al*., 2010; Galpaz *et al*., 2008; Kolotilin *et al*., 2007). Etiolated tissues from the Arabidopsis *crtiso* mutant (*ccr2*) accumulated an abundance of *cis*-carotenes; similar in amount to total cyclic carotenoid levels in WT (Cazzonelli *et al*., 2020). Leucoplasts may have an intrinsic property to hyperaccumulate *cis*-carotenes as evident in *tang*^*mic*^ roots where their abundance was 10.3-fold higher than the total of all cyclic and acyclic carotenoids quantified in WT roots. Acyclic carotenoids do not generally accumulate in WT plant tissues, but have been reported to occur in species- and tissue-dependent manner (Alagoz *et al*., 2018). Given that *cis*-carotenes were detected in WT MicroTom roots, either lower *CRTISO* expression and/or reduced CRTISO activity rate-limits the *cis*-to *trans* isomerisation of lycopene in roots. Therefore, light penetration to the root rhizosphere near the surface could mediate photoswitching between the *cis*-to *trans*-isoforms of lycopene and could perhaps affect the production of apocarotenoids that control root development.

### Roots of *tang*^*mic*^ show reduced ABA levels and mycorrhizal colonisation

Exogenous application of phytohormones has been shown to have reciprocal effects on AMF symbiotic interactions. Gibberellins and salicylic acid negatively regulate AMF symbiosis (Foo *et al*., 2013; Medina *et al*., 2003). The effect of cytokinin is not well understood, and the role of ethylene and jasmonic acid depends on their concentrations (Boivin *et al*., 2016; Khatabi and Schäfer, 2012; Ludwig-Müller *et al*., 2002). Positive regulators of the AMF symbiosis include auxin and ABA (Etemadi *et al*., 2014; Martín□Rodríguez *et al*., 2011). The *tang*^*mic*^ roots showed impaired *cis*/*trans*-lycopene isomerisation, and carotenoid profiling revealed that *tang*^*mic*^ roots did not accumulate neoxanthin or violaxanthin. Canonical biosynthesis of ABA proceeds via the NCED mediated oxidative cleavage of 9-*cis*-violaxanthin and 9-*cis*-neoxanthin; hence, there was a dramatic 23-fold reduction in ABA levels in *tang*^*mic*^ roots. ABA may also be synthesised by a non-canonical alternative, zeaxanthin epoxidase-independent pathway in plants (Jia *et al*., 2021a), but the 5-fold lower level of β-carotene in *tang*^*mic*^ roots indicated that this pathway is unlikely to generate ABA in the absence of neoxanthin or violaxanthin. The lack of ABA biosynthesis may be linked to the observed impairment in AMF colonisation, evident by a noticeable reduction in arbuscules and other AM structures compared to WT. While we attribute the reduction in ABA to the low AMF colonisation in *tang*^*mic*^ roots, we cannot rule out the possibility that a reduction in other apocarotenoids (e.g. zaxinone, strigolactone, and/or mycorradicin) could also contribute to the reduction of AM colonisation in *tang*^*mic*^ roots.

### *tang*^*mic*^ plants show reduced photosynthetic efficiency during drought stress

Soil water deficit conditions under drought trigger the accumulation of ABA in roots that moves through vascular tissues into foliar tissues where it elicits guard cell closure of stomata to limit transpiration and water loss, thereby maintaining leaf RWC (Bray, 2002; Kuromori *et al*., 2018; Sauter *et al*., 2001; Zhang *et al*., 2006). While the RWC of foliar tissues from WT and *tang*^*mic*^ consistently declined over a 15-day period of drought, there were significant differences in measurements of chlorophyll fluorescence between the genotypes. Particularly, the F_v_/F_m_ parameter remained mostly unchanged during the first phase of drought treatment in WT MicroTom foliar tissues, but the sudden deterioration in F_v_/F_m_ values observed in *tang*^*mic*^ leaf tissues. The latter effect is likely due to a lack of ABA signalling from the roots. Hence, there was a significant perturbation of PSII photochemistry or electron transport capacity within the *tang*^*mic*^ foliar photosystems that was irrespective of the initial decline in leaf RWC. This is in contrast to previous research where no clear relationship between the decline in RWC and F_v_/F_m_ was observed until the plant water reserves declined to critical levels (Woo *et al*., 2008). It is likely that ABA signalling from the roots is necessary to maintain maximum photosynthetic efficiency of PSII during the early stages of water deficit.

A very clear connection between the decline in the F_v_/F_m_ values and leaf RWC was identified for WT just prior to the terminal stage of drought stress (day 20) when the reserves reached a critical level. Surprisingly, a higher F_v_/F_m_ value and leaf RWC was maintained in *tang*^*mic*^ foliar tissues, and the plants survived for up to 30 days. One likely possible explanation for the extended survival of *tang*^*mic*^ is that because of its smaller shoot biomass and reduced root branching, the *tang*^*mic*^ plants would have had an overall lower requirement for water absorption within the rhizosphere, and in turn this could have prolonged the soil moisture content to enable a longer period of survival. Collectively, our findings revealed that the golden tangerine tomato variety might be drought sensitive, yet it can survive an extended period of drought due to having smaller shoot biomass.

### *tang*^*mic*^ plants show altered agronomic growth traits

The *tang*^*mic*^ plants display a bushy appearance due to increased shoot branching, similar to that displayed by CRTISO mutants in Arabidopsis (*ccr2*) and rice (*mit3*) where the latter showed enhanced tiller formation due to a SL deficiency (Cazzonelli *et al*., 2009; Dhami and Cazzonelli, 2020; Liu *et al*., 2018). The soil grown *tang*^*mic*^ plants displayed a noticeably smaller root system compared to WT, which was surprising given that the dry weight of root biomass was not different between WT and *tang*^*mic*^. When *tang*^*mic*^ roots were grown in MS medium, the root length and number of lateral roots were significantly reduced, agreeing with the root phenotypes observed in soil-grown plants. The increased root thickness of *tang*^*mic*^ may have compensated for the weight reduction caused by the smaller root architecture displayed by *tang*^*mic*^ plants. Rice loss-of-function *mhz5 CRTISO* mutant alleles also displayed shorter primary, adventitious, and lateral roots compared to wild-type seedlings (Yin *et al*., 2015). Furthermore, they had fewer adventitious roots but more lateral roots, indicating that impaired carotenoid isomerisation is required to promote important root agronomic traits (Yin *et al*., 2015). ABA accumulation was reduced in etiolated *mhz5* seedlings and *tang*^*mic*^ roots, yet the lack of ABA cannot be attributed to the reduction in primary root length. However, SLs have been reported to suppress adventitious root formation and reduce primary root length in Arabidopsis and pea (Rasmussen *et al*., 2012).

β-carotene derived apocarotenoids, anchorene and *iso*-anchorene promote anchor root development and inhibit primary root growth in Arabidopsis, respectively (Jia *et al*., 2019; Jia *et al*., 2021b). β-carotene derived β-cyclocitral promotes cell divisions in root apical meristem to enhance root growth development in rice, tomato and Arabidopsis (Dickinson *et al*., 2019). It is plausible that a lack of CAROTENOID ISOMERASE activity in *tang*^*mic*^ roots could affect the production of these apocarotenoid that have also been shown to regulate root development.

The *tang*^*mic*^ plants produced 50% less biomass, a consequence of having a decrease in plant height similar to what has been reported for rice CRTISO mutants, *zebra2-1* and *mit3*, that showed altered agronomic traits such as a decrease in plant height, seed setting and thousand-grain weight (Chai *et al*., 2011; Liu *et al*., 2018). *tang*^*mic*^ plants also showed a reduction in the numbers of flowers and fruit yield, indicating that the source (leaf) to sink (fruit) carbon allocation needed to sustain crop production was altered. The newly developed virescent leaves in *tang*^*mic*^ may have impaired photosynthesis. This is supported by observations made in the rice *zebra2-1* (*crtiso*) mutant which showed decreased photosynthetic rates and reduced chlorophyll content in yellow leaf sectors compared to green sectors that were still lower than that of the WT (Chai *et al*., 2011). In the rice *zel1* (*zebra-leaf1; crtiso*) mutant, the activity of photosynthetic oxygen evolution was also suppressed and accompanied by reduced LHCII trimer core proteins within the PS II supercomplexes and reduced plastoquinone (PQ) redox state (Wei *et al*., 2010). Hence, the smaller biomass phenotypes displayed by rice and tomato *crtiso* mutants are likely linked to reduced rates of photosynthesis in foliar tissues that disrupt the allocation of carbon for flower and fruit production. This could be one reason as to why the golden tangerine tomato has not become commercialised in protected cropping, despite being a highly abundant and nutritious source of prolycopene.

## CONCLUSION

The loss-of-function in CRTISO in the *tangerin*e MicroTom variety causes roots to accumulate *cis*-carotenes and block the biosynthesis of xanthophyll substrates required for ABA production. Photoisomerization of prolycopene can occur to some extent in root tissues but is rate-limited in the absence of any chlorophyll-derived photosensitizer. We conclude that the golden tomato variety *tang*^*mic*^ has reduced agronomical traits likely due to changes and/or the absence of carotenoid derivatives that control developmental processes. The poor mycorrhizal colonisation and drought responses in *tang*^*mic*^ might affect field crop performance, however, backyard or hydroponic glasshouse cultivation of golden heirloom varieties like *tangerine* could be a practical way to consume *cis-*isoforms of lycopene that can promote improved human health and prevent cancer and other inflammatory diseases. To make golden tomato varieties more attractive for commercialisation, golden varieties with reduced *CRTISO* gene expression specifically in the fruit would enable prolycopene accumulation yet not affect agronomic traits such as mycorrhizal colonisation in the rhizosphere.

## ACKNOWLEDGEMENT

We thank Mahfuza Pervin for helping to generate phenotypic measurements. We thank Meena Mikhael from Western Sydney University (WSU) for assistance with the hormone analyses at the WSU Mass Spectrometry Facility.

## AUTHOR CONTRIBUTIONS

CIC, JN and SA conceived ideas and designed the research. JN, SA and EF performed experiments and prepared figures. Data analysis involved all authors. JN wrote the manuscript with assistance from SA and primary supervisor CIC. Co-supervision of JN was provided by JMP and PK. Co-supervision of SA was provided by PK and ZC. All authors have read and edited the manuscript. The authors declare no conflict of interest.

